# Functionally conserved Pkd2, mutated in autosomal dominant polycystic kidney disease, localizes to the endoplasmic reticulum and regulates cytoplasmic calcium homeostasis in fission yeast

**DOI:** 10.1101/2022.09.20.508804

**Authors:** Takayuki Koyano, Kazunori Kume, Kaori Onishi, Makoto Matsuyama, Masaki Fukushima, Takashi Toda

## Abstract

Mutations in *PKD1* or *PKD2* genes lead to autosomal dominant polycystic kidney disease (ADPKD) that is the most frequent family inherited renal disorder. These genes encode polycystin-1/PC-1 and polycycstin-2/PC-2, respectively. Although the genetic basis of ADPKD is well established, the crucial functions of polycystins underlying onset and development of cyst formation remain elusive. Fission yeast *Schizosaccharomyces pombe* has a single polycystin homolog, Pkd2, which is essential for cell growth. In this study, the truncation analyses of Pkd2 reveal that Pkd2 localizes to not only the plasma membrane but also the endoplasmic reticulum (ER) and regulates cytoplasmic calcium signaling in fission yeast. Internal transmembrane domains within Pkd2 are sufficient for these processes. Surprisingly, more than half of Pkd2 is not required for cell viability. Cytoplasmic calcium levels are mainly regulated through C-terminus of Pkd2. Importantly, human Pkd2 also localizes to the ER and furthermore, fully complements the loss of fission yeast Pkd2. As the functions of polycystin-2 are conserved, fission yeast provides a suitable model to study the mechanism of ADPKD as well as polycystins.

## Introduction

Autosomal dominant polycystic kidney disease (ADPKD) is one of the most frequent genetic inherited renal diseases in the world, with an estimated prevalence of one in 1,000 individuals (Lanktree et al., 2018). The disease responsible genes, *PKD1* and *PKD2* which encode polycystin-1 (PC-1) and polycystin-2 (PC-2) respectively, have already been identified (Hughes et al., 1995; Mochizuki et al., 1996). PC-1 consists of a large N-terminal extracellular domains, 11 transmembrane domains and a C-terminal cytoplasmic region (Ong and Harris, 2015). PC-2, a member of the transient receptor potential (TRP) superfamily, has six transmembrane regions and a C-terminal cytoplasmic region. The C-terminus contains an EF-hand motif, a coiled-coil domain, and an endoplasmic reticulum (ER) retention motif (Brill and Ehrlich, 2020; Cai et al., 1999; Vien et al., 2020), whereas the N-terminus possess cilia localization sequence (Geng et al., 2006). It has been suggested that PC-1 and PC-2 function in the same pathway, as mutations in either gene cause the similar cyst phenotypes (Harris and Torres, 2014). In agreement with this, initial studies suggest that PC-2 interacts with PC-1 via each C-terminus and localizes to cilia in a PC-1 dependent manner (Grieben et al., 2017; Hanaoka et al., 2000; Ong and Harris, 2015). The mechanism by which loss of ciliary PC-1 or PC-2 function results in cyst formation through decreasing in cytoplasmic calcium levels and increase in cAMP is thought to be central to ADPKD pathology for a long time (Cornec-Le Gall et al., 2019; Torres and Harris, 2014). In addition, structural analysis revealed that human PC-1 and PC-2 form a hetero-tetramer in 1:3 ratio (Su et al., 2018). However, it was also reported that PC-2 predominantly localized to the ER, forming a homo tetramer, acting as a cation channel independent of PC-1 (Cai et al., 1999; Liu et al., 2018; Shen et al., 2016). In addition, recent study suggested that ER localized PC-2 contributes to calcium release and anti-cystogenesis (Padhy et al., 2022). Thus, ADPKD pathology associated functions of polycystins remain to be established.

Fission yeast *Schizosaccharomyces pombe*, a unicellular model organism, has Pkd2 which is essential for cell viability, as a solo homolog of polycystins. Pkd2 consists of the N-terminal extracellular region, the nine transmembrane domains and the C-terminal cytoplasmic region. The roles of each region, however, have not been determined yet. Pkd2 reportedly localizes to the plasma membrane and cell division site (septum in fungi) (Morris et al., 2019). Mutations in *pkd2* cause cell separation defect and fails to maintain cellular integrity (Morris et al., 2019; Sinha et al., 2022). Furthermore, overexpression of *pkd2* causes growth arrest and increase of cytoplasmic calcium levels (Ma et al., 2011; Palmer et al., 2005).

Here, we carried out the truncation analyses of fission yeast Pkd2 and revisited its cellular localization. We show that Pkd2 localizes to the ER rather than the septum site and regulates the cytoplasmic calcium homeostasis. More than a half of N-terminus is not required for cell viability. C-terminus, including the latter four transmembrane regions has a central role in the regulation of cytoplasmic calcium homeostasis. Interestingly, human *PKD2* fully complements the loss of *pkd2* in fission yeast.

## Results and Discussion

### 1. The transmembrane region of Pkd2 is required for cell survival and cytoplasmic calcium homeostasis

Previous reports suggested that overexpression of *pkd2* results in cell death accompanied with increased calcium levels in the cytoplasm (Ma et al., 2011; Palmer et al., 2005). In order to analyze the requirement of each region of Pkd2 for the growth and calcium signaling, we constructed and overexpressed a series of truncations in fission yeast cells (Fig. 1A). The CDRE (Calcineurin Dependent Response Element) reporter system was used to estimate the cytoplasmic calcium level (Kume et al., 2017; Kume et al., 2011). As expected, the CDRE-dependent transcription was increased by adding CaCl_2_ in a Prz1-dependent manner (Fig. 1B). Neither deletion of N-nor C-terminus of Pkd2 affected the growth and CDRE activation (Fig. 1C, D). N-or C-terminus of Pkd2 alone did not affect either. Interestingly, the internal transmembrane region was sufficient to induce both growth inhibition and CDRE activation (Fig. 1C, D). These data suggest that the transmembrane region plays a critical role for Pkd2, while either N-or C-terminal region is dispensable.

**Figure 1.**
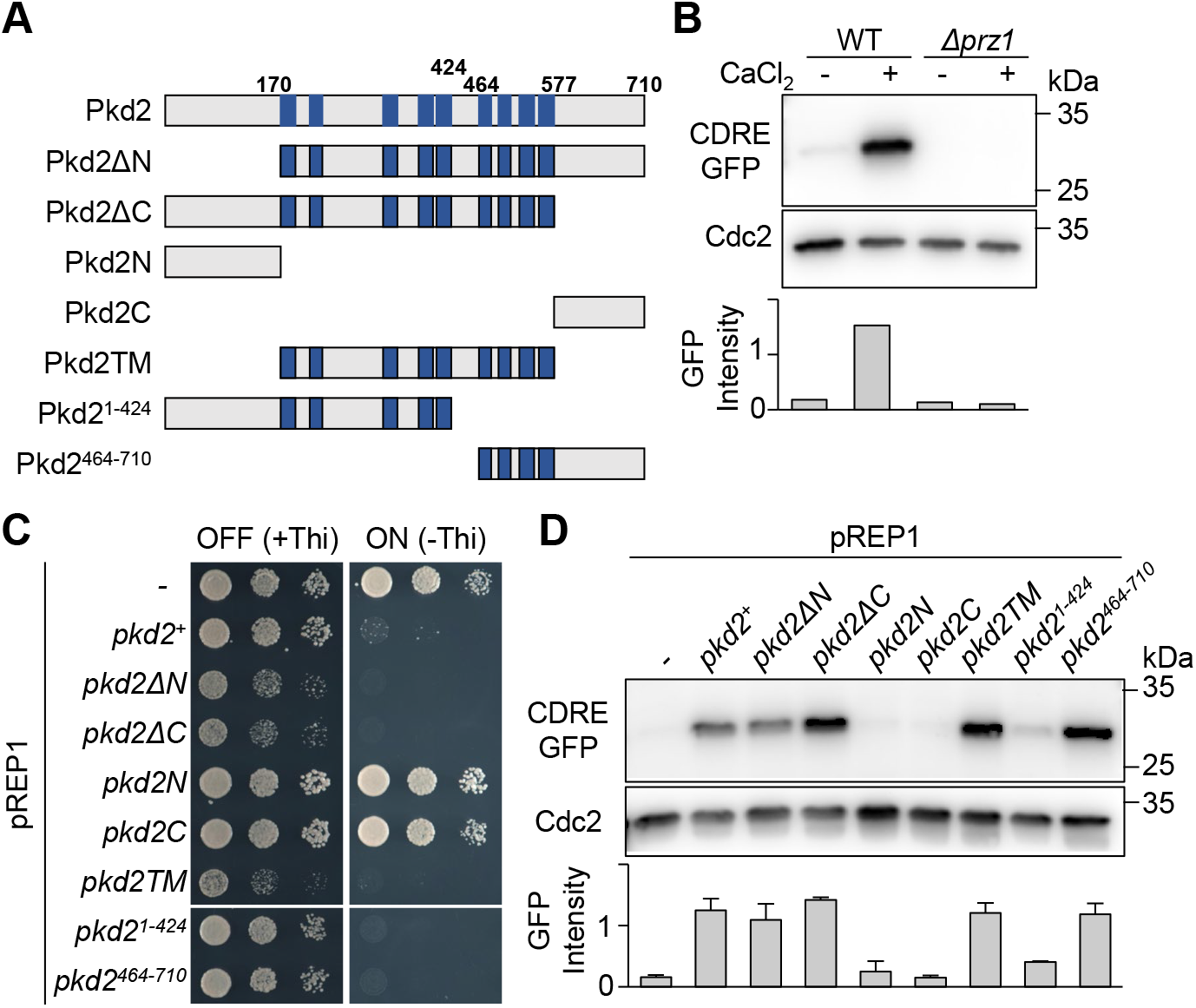
Transmembrane region of Pkd2 is required for the functions. (A) Schematic structures of Pkd2 truncations. Dark blue boxes indicate transmembrane (TM) region. (B) Calcineurin activity determined by CDRE-GFP reporter system. Indicated strains were cultured in YE5S at 27ºC with or without 0.1 M CaCl_2_ for 2 h. Calcineurin activity was estimated by immunoblotting with anti-GFP antibodies. (C) Spot tests. Indicated strains were serially diluted and spotted onto PMG with or without thiamine. The plates were kept at 27ºC for 4 days. (D) Calcineurin activity determined by CDRE-GFP reporter system. Indicated strains were cultured in PMG without thiamine for 18 h to induce overexpression. Calcineurin activity was estimated by immunoblotting with anti-GFP antibodies.

To classify the roles of the transmembrane domains, we divided Pkd2 into Pkd2^1-424^, which includes the first five transmembrane domains and Pkd2^464-710^, which includes the latter four transmembrane domains. Overexpression of either construct inhibited the growth like full length (Fig. 1C), indicating that each of them is capable of inducing cytotoxicity upon overexpression. With regards to calcium signaling, the overexpression of *pkd2*^*1-424*^ failed to increase cytoplasmic calcium levels, whereas the overexpression of *pkd2*^*464-710*^ showed noticeable increase, comparable to full length (Fig. 1D). These data suggest that each of the first five or the latter four transmembrane regions has an independent role for the growth, and that C-terminus including the latter four transmembrane regions regulates the cytoplasmic calcium homeostasis.

### 2. Pkd2 localizes to the ER and the internal transmembrane region is sufficient for its localization

Previous work showed that Pkd2 localizes to the plasma membrane, concentrated at the cell tip and the medial cell division site/septum (Morris et al., 2019). This localization pattern was obtained using C-terminally tagged GFP (Pkd2-GFP). We checked the localization of full-length Pkd2 and its truncations that contained N-terminally tagged GFP (GFP-Pkd2) from the plasmid in the cells. Intriguingly, full length of Pkd2 (GFP-Pkd2) colocalized with Pmr1, a marker for the ER (Nakazawa et al., 2019), indicating that Pkd2 may localize to the ER (Fig. 2A). Localization of Pkd2ΔN170, Pkd2ΔC577, or Pkd2TM (Fig. 2B) was identical to full length Pkd2 (Figs 2B, S1A); no signals were evident at the septum. N-terminus of Pkd2 (Pkd2N) localized to the cytoplasm and weakly to the nuclear periphery, whereas C-terminus of Pkd2 (Pkd2C) displayed a uniform cytoplasmic pattern (Fig. 2B, S1A). These data suggest that the transmembrane region is sufficient for ER localization. We noted that although GFP-Pkd2 and various truncations were expressed from plasmids under the *nmt41* prompter (Fig. S1B), this level of expression did not compromise the growth even under the inducible condition (Fig. S1C).

**Figure 2.**
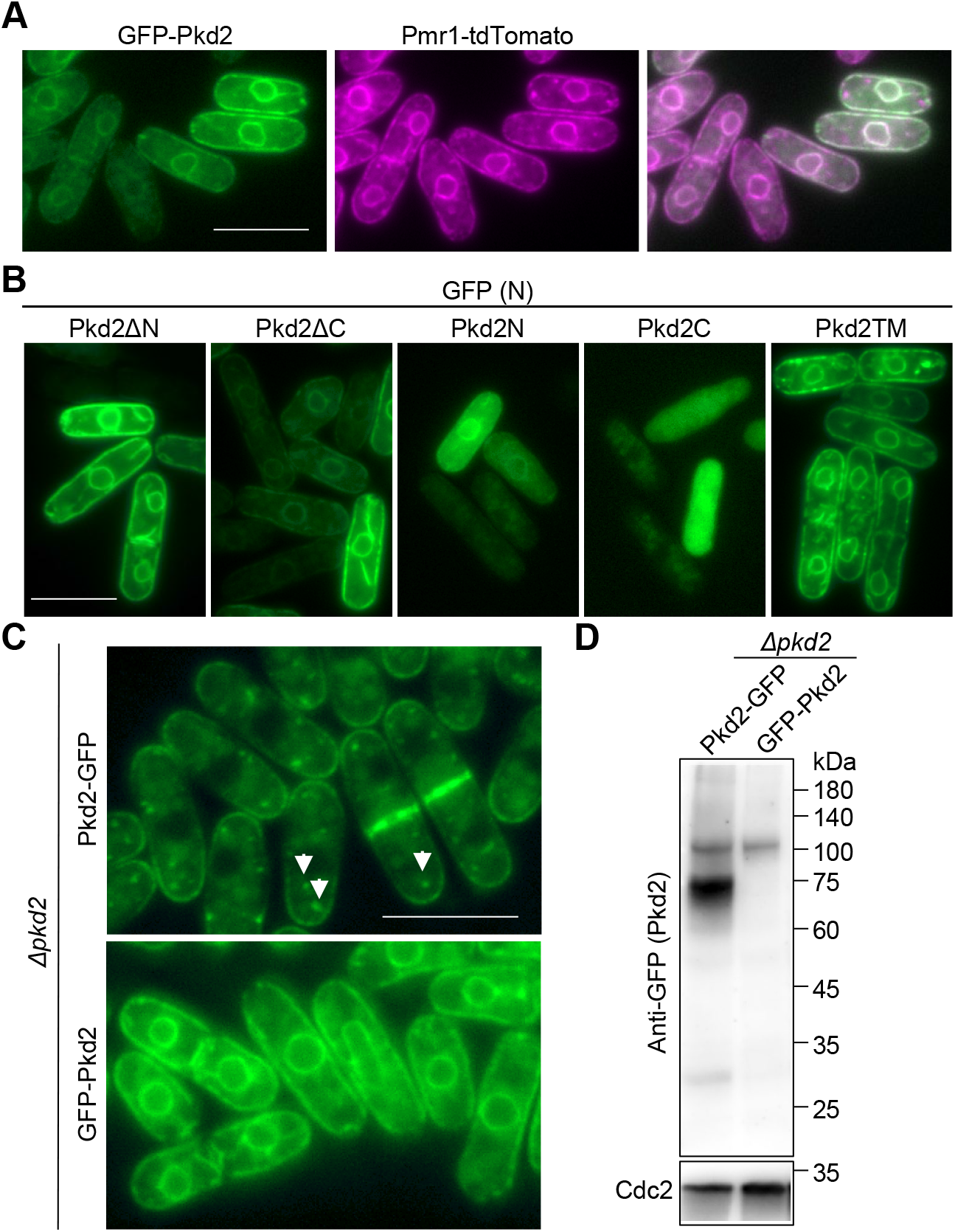
N-terminally tagged Pkd2 localizes to the ER. (A, B) Localization of GFP-Pkd2 (A) or GFP-Pkd2 truncations (B) expressed from plasmids in Pmr1-tdTomato expressing cells. For truncations, only GFP images were shown. Pmr1-tdTomato and merged images were shown in supplementary Figure S1A. Bars; 10 μm. (C) Localization of Pkd2. Pkd2-GFP or GFP-Pkd2 expressed from its own promoter in *Δpkd2* background. (D) Whole cell extracts were prepared from the indicated strains and immunoblotting carried out with anti-GFP and anti-Cdc2 (as a control) antibodies. The positions of size markers are shown on the right.

To decipher whether Pkd2’s ER localization is due to overexpression, we integrated a single copy of *GFP-pkd2* or *pkd2-GFP* under the endogenous promoter in the *pkd2* deleted background. As previously reported (Morris et al., 2019), C-terminally tagged Pkd2 (Pkd2-GFP) localized to the septum and the plasma membrane, especially enriched at cell tips (Fig. 2C). It is noted that Pkd2-GFP signals were also detected as numerous cytoplasmic dots (Fig. 2C, arrowheads). On the other hand, N-terminally tagged Pkd2 (GFP-Pkd2) localized to the ER, like a plasmid-derived expression (Fig. 2A, C). The protein expression of both GFP-Pkd2 and Pkd2-GFP was examined by immunoblotting analysis (Fig. 2D). GFP-Pkd2 showed only a single band around the expected full-length size, whereas Pkd2-GFP expressing cells showed two prominent bands: expected full-length size (∼110 kDa) and smaller size (∼75 kDa). Overall, GFP tagging of Pkd2 to its C-terminus resulted in different localization patterns and susceptibility to proteolytic cleavage. Given the results of GFP-Pkd2 that did not lead to the emergence of proteolysis or cytoplasmic dots, we posit that fission yeast Pkd2 localizes to the ER via the transmembrane regions.

Why C-terminally-GFP tagged Pkd2 (Pkd2-GFP) localizes to the plasma membrane is currently unknown. It is possible that C-terminal tagging affects the Pkd2 expression level and the localization, as the different sized band was included in the extract from Pkd2-GFP expressing cells. Polycycstin-2 levels are regulated by microRNAs (miRNAs) and contribute to cyst formation (Lee et al., 2019). Recent study suggests that the deletion of miR-17 binding sites within the 3’UTR increased expression levels of polycystin-2 and attenuated the cyst formation in mouse ADPKD model (Lakhia et al., 2022). In our C-terminal tagging constructs, its own 3’UTR regions are inactivated. It is interesting whether 3’UTR region of fission yeast Pkd2 also contributes to their expression and miRNA cis-inhibition system is evolutionary conserved.

### 3. The C-terminal region of Pkd2 plays a central role in cytoplasmic calcium regulation

We then constructed several truncations within GFP-Pkd2 that were expressed from its own promoter in the absence of endogenous *pkd2* (Fig. 3A). N-terminal deletions up to 170 amino acid residues did not affect either viability or ER localization (Fig. 3B). Remarkably, even a further truncation construct, GFP-Pkd2ΔN424 expressing cells, in which the first five transmembrane regions are deleted, was viable and localized to the ER like full length GFP-Pkd2 (Fig. 3B). Although Pkd2 has a signal sequence in N-terminus (Morris et al., 2019), the N-terminal deletions did not affect to the ER localization, suggesting that Pkd2 localizes to the ER via multiple signals. On the other hand, a C-terminal deletion, GFP-Pkd2ΔC577 was viable, but displayed noticeable defects: localization to the cortical ER was weakened and cells showed abnormal morphologies including multiseptation (Fig. 3B). The protein levels were examined by immunoblotting analyses, and we found that the band signals of GFP-Pkd2ΔN170 and GFP-Pkd2ΔC577 were somewhat weakened (Figure 3C). In addition, the C-terminal deletion strain was hypersensitive to CaCl_2_ (Figure 3D), indicating that the C-terminal region is required for cytoplasmic calcium homeostasis. Taking these data together, we concluded that each of the first five and the latter four transmembrane regions has a mutually complementing activity for cell viability and that C-terminus plays a major role in ER localization and calcium homeostasis. Other groups studies also suggest that fission yeast Pkd2 appears to regulate the cytoplasmic calcium levels (Ma et al., 2011). We envisage that the C-terminal region including the latter four transmembrane domains has a crucial role in calcium regulation, as the deletion of the C-terminal region caused more severe defects and C-terminus overexpression was sufficient to increase the cytoplasmic calcium levels. N-terminus is not required for the cell viability, however, the overexpression of N-terminus leads to the cell death as well as C-terminus. It appears that Pkd2 has two independent functions, but somehow compensates with each other. Further analysis, such as the identification of the molecules which genetically and/or physically interact with Pkd2 will uncover the importance of each region.

**Figure 3.**
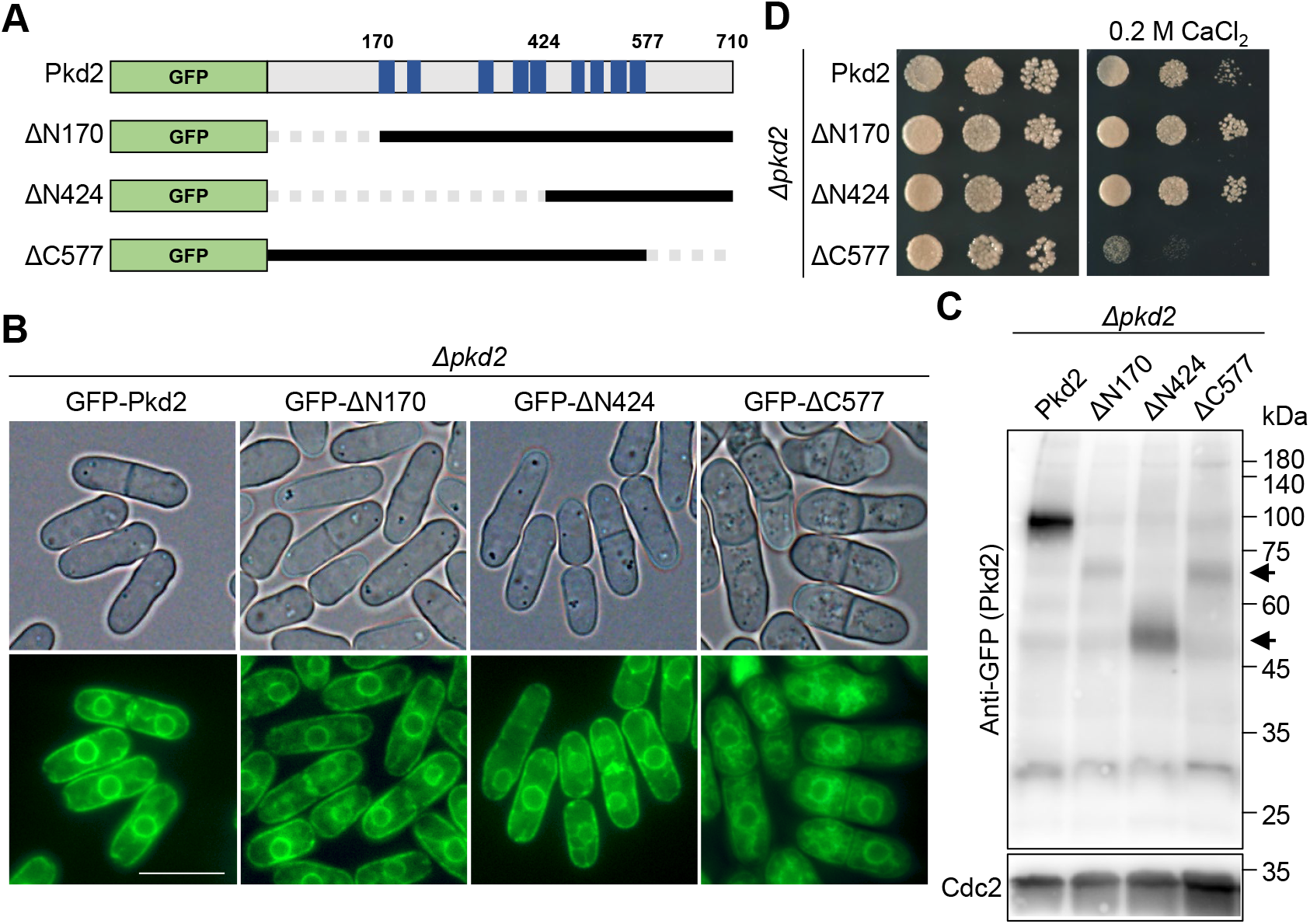
The C-terminus of Pkd2 is required for the viability and cytoplasmic calcium homeostasis. (A) Schematics of truncations. Gray-dashed lines indicated deleted regions. (B) Localization of GFP-Pkd2 truncations expressed from its own promoter in *Δpkd2* background. Bar; 10 μm. (C) Whole cell extracts were prepared from the indicated strains and immunoblotting carried out with anti-GFP and anti-Cdc2 (as a control) antibodies. Arrowheads indicate the position of the expected bands. The positions of size markers are shown on the right. (D) Spot tests. Serially diluted strains were spotted onto YE5S with or without 0.2 M CaCl_2_ and incubated at 27ºC for 3days.

### 4. Human *PKD2* complements fission yeast *pkd2*

To examine functional complementation of Pkd2 between humans and fission yeast, we expressed human *PKD2* (*hpkd2*) in the absence of fission yeast Pkd2. Cells expressing GFP-hPkd2 or hPkd2-GFP under the *pkd2* promoter in *Δpkd2* background were viable even under the high CaCl_2_ condition (Fig. 4A), suggesting that human *PKD2* can effectively complement the loss of fission yeast *pkd2*. Both GFP-hPkd2 and hPkd2-GFP localized to the ER (Fig. 4B), and the expected sized proteins (∼140 kDa) were expressed in fission yeast (Fig. 4C), indicating that the functional proteins were produced. We noted that the existence of cleavage products (∼75 kDa) in hPkd2-GFP-expressing cells, which is similar to that produced from fission yeast Pkd2-GFP, though its amount was substantially lower. We conclude that *PKD2* genes are evolutionary conserved between humans and fission yeast.

**Figure 4.**
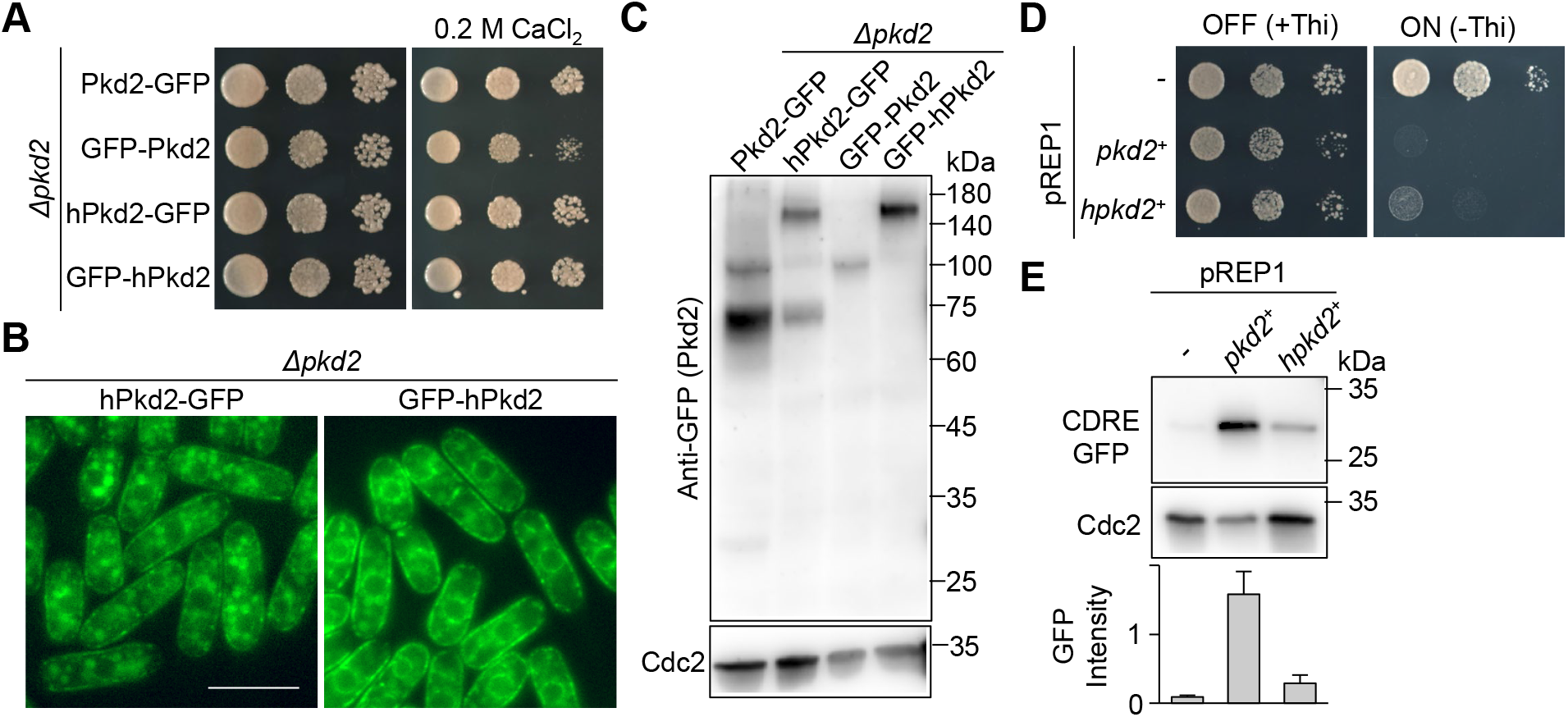
Human *PKD2* complements fission yeast *pkd2*. (A) Spot tests. Serially diluted strains were spotted onto YE5S with or without 0.2 M CaCl_2_ and incubated at 27ºC for 3days. (B) Localization of hPkd2-GFP or GFP-hPkd2 expressed from *pkd2* promoter in *Δpkd2* cells. Bar; 10 μm. (C) Protein levels of hPkd2. Whole cell extracts were prepared from the indicated strains and immunoblotting carried out with anti-GFP and anti-Cdc2 (as a control) antibodies. The positions of size markers are shown on the right. (D) Spot tests. Indicated strains were spotted onto PMG with or without thiamine and incubated at 27ºC for 4 days. (E) Calcineurin activity. Indicated strains were cultured in PMG without thiamine for 18 h to induce overexpression. Calcineurin activity was estimated by immunoblotting with anti-GFP antibodies.

We next overexpressed *hpkd2* in fission yeast. Overexpression of *hpkd2* also activated cytoplasmic calcium and caused growth arrest in fission yeast (Fig. 4D, E); therefore, human Pkd2 recapitulated the physiological impact caused by fission yeast Pkd2. These data reinforce the notion that human polycystin-2 is functional in fission yeast and has similar functions to Pkd2. The level of calcineurin activation, however, was not as much as that of fission yeast *pkd2* overexpression (Figure 4E), indicating that hPkd2 might not fully substitute for cellular physiologies derived from fission yeast Pkd2.

Recent work suggests that polycystin-2 is a non-selective cation channel and involved in permeability of other monovalent cations (Liu et al., 2018; Shen et al., 2016). Consistent with this, the effect of *hpkd2* overexpression on cytoplasmic calcium levels was limited in this study. Interestingly, the overexpression effect of N-terminal region (*pkd2*^*1-424*^) on calcium homeostasis is also minor. One possibility is that the N-terminal region is conserved in human and fission yeast, and C-terminus of fission yeast Pkd2 is an additional, divergent region. Indeed, human Pkd2 consists of six transmembrane domains, whereas fission yeast has nine. Furthermore, human Pkd2 have a large extracellular polycystin domain (∼200 amino acid long; also referred TOP domain), in which ADPKD pathogenic gene mutations are accumulated, and it contributes to channel assembly and function (Douguet et al., 2019; Grieben et al., 2017; Shen et al., 2016; Wilkes et al., 2017). This polycystin domain exists like a physical ‘lid’ of the channel (Douguet et al., 2019; Shen et al., 2016). Fission yeast Pkd2 has a large extracellular region in N-terminus (1-170 amino acid). So far, what N-terminus does in the context of cellular physiology is unclear, but at least the overexpression causes cell death. Conservation of the functional and structural polycystin domain is of great interest and its roles should be investigated in the future.

Nonetheless, human *PKD2* complements the loss of *pkd2* in fission yeast and its gene protein like that of fission yeast localizes to the ER; hPkd2 appears to be functionally conserved. Thus, our data provide a potential that fission yeast is an excellent model organism in which to study the molecular basis of ADPKD as well as the cellular function(s) of Pkd2.

## Materials and Methods

### Yeast general method

Standard media and methods for fission yeast were used. C-terminal tagging and gene deletion were carried out with the PCR-based method using homologous recombination (Bahler et al., 1998; Sato et al., 2005). The full length of *pkd2* gene or *pkd2-GFP-hphMX*, including 5’UTR region (324 bases) and 3’UTR region (569 bases) were amplified from the genomic DNA of wild type or *pkd2-GFP-hphMX* strain, respectively. The genomic DNA were prepared with GenTLE (Takara Bio, Japan). The fragments were subcloned into the integrated plasmid pJK148 carrying the *leu1*^*+*^ gene, the resulting plasmids were linearized by *Nru*? within the *leu1*^*+*^ gene and integrated to the *leu1-32* locus. For truncations and N-terminal GFP tagged strains construction, the fragments were amplified with PrimeSTAR MAX (Takara Bio, Japan, R045) with the appropriate primer pairs having overlapping sequence. The purified fragments were integrated into digested vectors by using In-fusion Snap Assembly Master mix (Takara Bio, Japan, 638947). Our lab stock plasmids, pREP1, pREP41-GFP(N), and pJK148 were used as the vector. Human *PKD2* cDNA was purchased (Horizon Discovery, MHS6278-211688893). Multiple tagging strains were constructed by random spore. Strains used in this study were listed in supplementary table S1. Strains were grown in rich YE5S media and incubated at 27 °C unless otherwise stated. PMG or EMM media (Sunrise Science Products, TN, U.S.A., 2060 or 2005) were used to induce overexpression. Appropriate supplements (leucine, lysine, histidine, uracil, and adenine) were added at concentration of 50 mg/L each if required.

### Microscope

Fluorescent microscope images were obtained by the Olympus IX83 inverted microscope system with UPLXAPO 60x objective lens (NA 1.42, immersion oil) and a DP80 digital camera. Cells were collected by the centrifuge at 5,000rpm for 1 min, and spotted onto glass slide. The cells were observed immediately after coverslip. Images were processed and analyzed by using CellSens Dimension (OLYMPUS, Japan) and Adobe PhotoShop.

### Western blotting

Whole cell extracts were prepared according to the alkaline method (Matsuo et al., 2006). 10-20 ml of cell cultures (OD_600_ : 0.4-0.7) were centrifuge at 3,500 rpm for 1 min. The cells were washed with 1 ml of distilled water, centrifuged for 1 min. After resuspending in 0.3 M NaOH, the cells were kept at room temperature for 10 min with shaking. Discarding the supernatant, sample buffer (60 mM Tris-HCl (pH6.8), 4% β-Mercaptoethanol, 4% SDS, 0.01% BPB, 5% glycerol) was added, and the samples stayed at room temperature for at least 1 h. The samples were separated by 10% of SDS-PAGE gel (Bio-rad, CA, U.S.A., 4561035) and transfer to polyvinylidene difluoride membrane. The membranes were blocked with 5 % of skimmed milk in TBS-tween20 (TBST) for 30 min at room temperature, subsequently incubated with primary antibodies diluted in TBST at 4ºC for overnight. Anti-GFP (MBL, Japan, 598) and Anti-GFP (Roche, U.S.A., 11814460001) were used for CDRE-GFP, GFP tagged Pkd2, respectively. After washing, the membranes were incubated with appropriate secondary antibodies at room temperature for 60 min; anti-Rabbit (Thermo Fisher Scientific, G-21234), anti-Mouse (Thermo fisher Scientific, G-21040). Then the membranes were incubated with Clarity (Bio-Rad, 1705061) or Clarity Max (Bio-Rad, 1705062) western ECL substrate. For the control, the membranes were re-incubated with anti-Cdc2 (SantaCruz Biotechnology, Texas, U.S.A., SC-53217) in TBST with 0.1% of sodium azide at room temperature for 3 h. Amersham Image Quant 800 (Cytiva, Japan) was used for detection of chemiluminescence. The band intensities were measured by ImageJ. The CDRE-GFP intensities were normalized by dividing by Cdc2 intensities. The bar graphs showed the average of three experiments, with exception of Figure 1B. Error bars indicated standard deviation (S.D.).

## Acknowledgements

We thank Dr Fumihiro Shigei, the Chairman of the Board, Sowa-kai Medical Foundation, for encouragement and financial support.

## Supplementary data

**Supplementary table S1.** Strains in this study.

|  | Genotype |
| --- | --- |
| TK1042-1 | *h- ura4-294::CDRE:GFP leu1-32* |
| TK1230-1 | *h- Δprz1::kanMX ura4-294::CDRE:GFP leu1-32* |
| TK1223 | *h- ura4-294::CDRE:GFP leu1-32/pREP1 (Transformant)* |
| TK1224 | *h- ura4-294::CDRE:GFP leu1-32/pREP1-pkd2 (Transformant)* |
| TK1242 | *h- ura4-294::CDRE:GFP leu1-32/pREP1-pkd2ΔN170 (Transformant)* |
| TK1255 | *h- ura4-294::CDRE:GFP leu1-32/pREP1-pkd2Δ577 (Transformant)* |
| TK1265 | *h- ura4-294::CDRE:GFP leu1-32/pREP1-pkd2N (Transformant)* |
| TK1286 | *h- ura4-294::CDRE:GFP leu1-32/pREP1-pkd2C (Transformant)* |
| TK1266 | *h- ura4-294::CDRE:GFP leu1-32/pREP1-pkd2TM (Transformant)* |
| TK1445 | *h- ura4-294::CDRE:GFP leu1-32/pREP1-pkd2<sup>1-424</sup> (Transformant)* |
| TK1446 | *h- ura4-294::CDRE:GFP leu1-32/pREP1-pkd2<sup>465-710</sup> (Transformant)* |
| TK1225 | *h- ura4-294::CDRE:GFP leu1-32/pREP1-hpkd2 (Transformant)* |
| TK1396 | *h- pmr1<sup>+</sup>:tdTomato:natMX leu1-32/pREP41-GFP(N)-pkd2 ura4-D18 (Transformant)* |
| TK1397 | *h- pmr1<sup>+</sup>:tdTomato:natMX leu1-32/pREP41-GFP(N)-pkd2ΔC577 ura4-D18 (Transformant)* |
| TK1398 | *h- pmr1<sup>+</sup>:tdTomato:natMX leu1-32/pREP41-GFP(N)-pkd2ΔN170 ura4-D18 (Transformant)* |
| TK1399 | *h- pmr1<sup>+</sup>:tdTomato:natMX leu1-32/pREP41-GFP(N)-pkd2N ura4-D18 (Transformant)* |
| TK1400 | *h- pmr1<sup>+</sup>:tdTomato:natMX leu1-32/pREP41-GFP(N)-pkd2TM ura4-D18 (Transformant)* |
| TK1390 | *h- pmr1<sup>+</sup>:tdTomato:natMX leu1-32/pREP41-GFP(N)-pkd2C ura4-D18 (Transformant)* |
| TK1129-1 | *h- Δpkd2::kanMX leu1-32::pkd2<sup>+</sup>:GFP:hphMX (integrated)* |
| TK1323-1 | *h- Δpkd2::kanMX leu1-32::P<sub>pkd2</sub>-GFP-pkd2<sup>+</sup>-T<sub>pkd2</sub> (integrated)* |
| TK1408-1 | *h+ Δpkd2::kanMX leu1-32::P<sub>pkd2</sub>-GFP-pkd2ΔN170-T<sub>pkd2</sub> (integrated) his2* |
| TK1388-1 | *h- Δpkd2::kanMX leu1-32::P<sub>pkd2</sub>-GFP-pkd2ΔN424-T<sub>pkd2</sub> (integrated) ura4-D18* |
| TK1455-1 | *h- Δpkd2::kanMX leu1-32::P<sub>pkd2</sub>-GFP-pkd2Δ577-T<sub>pkd2</sub> (integrated)* |
| TK1375-1 | *h- Δpkd2::kanMX leu1-32::hpkd2<sup>+</sup>:GFP:hphMX (integrated)* |
| TK1401-1 | *h- Δpkd2::kanMX leu1-32::P<sub>pkd2</sub>-GFP-hpkd2<sup>+</sup>-T<sub>pkd2</sub> (integrated)* |

**Figure S1.**
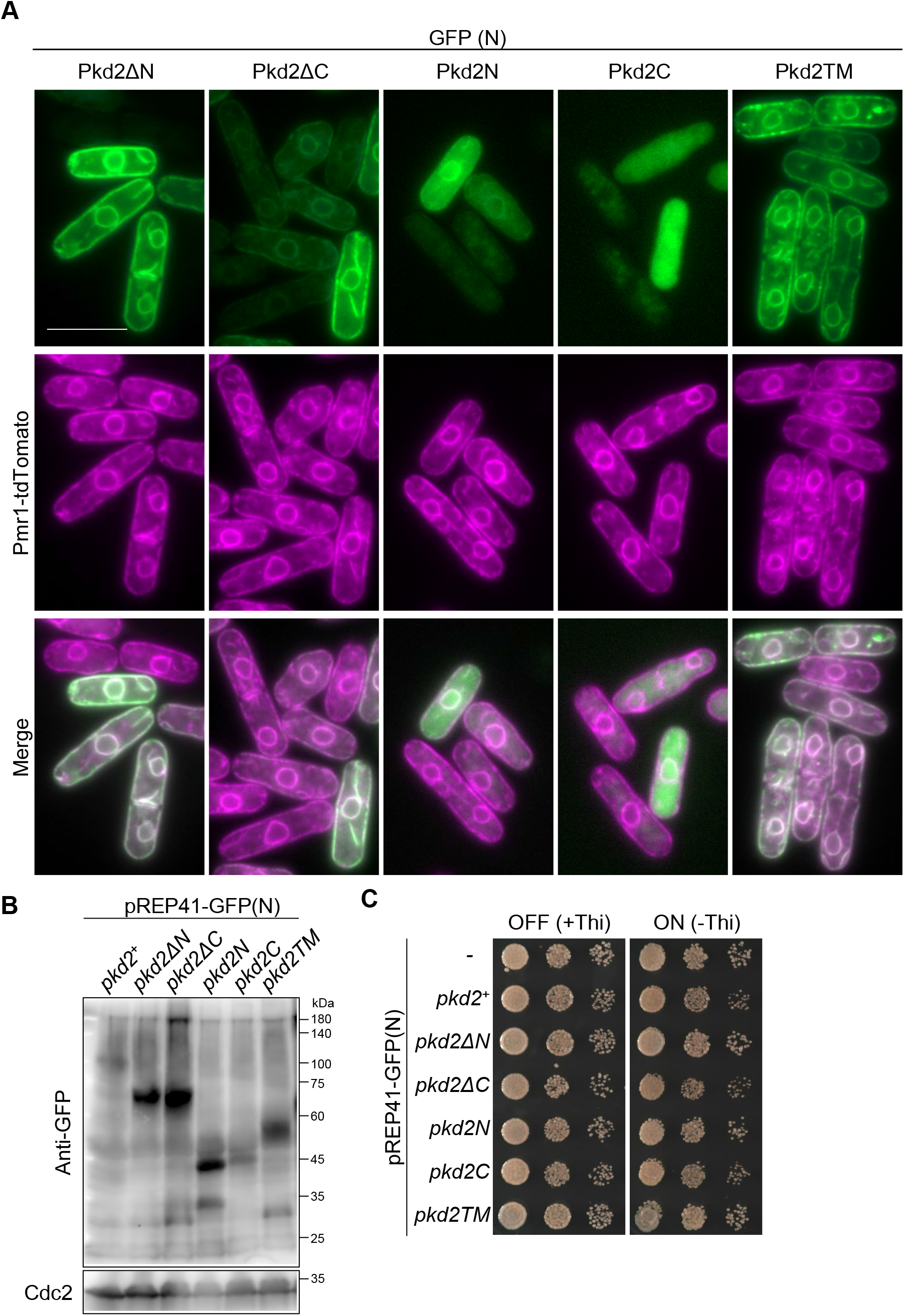
N-terminal GFP tagged Pkd2 localize to ER. (A) Localization of GFP-Pkd2 or GFP-Pkd2 truncations. Identical GFP images are shown in Fig. 2B. (B) Whole cell extracts were prepared from the indicated strains and immunoblotting carried out with anti-GFP and anti-Cdc2 antibodies. (C) Spot test. Spot-test. Indicated strains were serially diluted and spotted onto EMM with or without thiamine. The plates were kept at 27ºC for 3 days.

## References

Bahler, J., Wu, J. Q., Longtine, M. S., Shah, N. G., McKenzie, A., 3rd, Steever, A. B., Wach, A., Philippsen, P. and Pringle, J. R. (1998). Heterologous modules for efficient and versatile PCR-based gene targeting in Schizosaccharomyces pombe. Yeast 14, 943–51.

Brill, A. L. and Ehrlich, B. E. (2020). Polycystin 2: A calcium channel, channel partner, and regulator of calcium homeostasis in ADPKD. Cell Signal 66, 109490.

Cai, Y., Maeda, Y., Cedzich, A., Torres, V. E., Wu, G., Hayashi, T., Mochizuki, T., Park, J. H., Witzgall, R. and Somlo, S. (1999). Identification and characterization of polycystin-2, the PKD2 gene product. J Biol Chem 274, 28557–65.

Cornec-Le Gall, E., Alam, A. and Perrone, R. D. (2019). Autosomal dominant polycystic kidney disease. Lancet 393, 919–935.

Douguet, D., Patel, A. and Honore, E. (2019). Structure and function of polycystins: insights into polycystic kidney disease. Nat Rev Nephrol 15, 412–422.

Geng, L., Okuhara, D., Yu, Z., Tian, X., Cai, Y., Shibazaki, S. and Somlo, S. (2006). Polycystin-2 traffics to cilia independently of polycystin-1 by using an N-terminal RVxP motif. J Cell Sci 119, 1383–95.

Grieben, M., Pike, A. C., Shintre, C. A., Venturi, E., El-Ajouz, S., Tessitore, A., Shrestha, L., Mukhopadhyay, S., Mahajan, P., Chalk, R. et al. (2017). Structure of the polycystic kidney disease TRP channel Polycystin-2 (PC2). Nat Struct Mol Biol 24, 114–122.

Hanaoka, K., Qian, F., Boletta, A., Bhunia, A. K., Piontek, K., Tsiokas, L., Sukhatme, V. P., Guggino, W. B. and Germino, G. G. (2000). Co-assembly of polycystin-1 and -2 produces unique cation-permeable currents. Nature 408, 990–4.

Harris, P. C. and Torres, V. E. (2014). Genetic mechanisms and signaling pathways in autosomal dominant polycystic kidney disease. J Clin Invest 124, 2315–24.

Hughes, J., Ward, C. J., Peral, B., Aspinwall, R., Clark, K., San Millan, J. L., Gamble, V. and Harris, P. C. (1995). The polycystic kidney disease 1 (PKD1) gene encodes a novel protein with multiple cell recognition domains. Nat Genet 10, 151–60.

Kume, K., Hashimoto, T., Suzuki, M., Mizunuma, M., Toda, T. and Hirata, D. (2017). Identification of three signaling molecules required for calcineurin-dependent monopolar growth induced by the DNA replication checkpoint in fission yeast. Biochem Biophys Res Commun.

Kume, K., Koyano, T., Kanai, M., Toda, T. and Hirata, D. (2011). Calcineurin ensures a link between the DNA replication checkpoint and microtubule-dependent polarized growth. Nat Cell Biol 13, 234–42.

Lakhia, R., Ramalingam, H., Chang, C. M., Cobo-Stark, P., Biggers, L., Flaten, A., Alvarez, J., Valencia, T., Wallace, D. P., Lee, E. C. et al. (2022). PKD1 and PKD2 mRNA cis-inhibition drives polycystic kidney disease progression. Nat Commun 13, 4765.

Lanktree, M. B., Haghighi, A., Guiard, E., Iliuta, I. A., Song, X., Harris, P. C., Paterson, A. D. and Pei, Y. (2018). Prevalence Estimates of Polycystic Kidney and Liver Disease by Population Sequencing. J Am Soc Nephrol 29, 2593–2600.

Lee, E. C., Valencia, T., Allerson, C., Schairer, A., Flaten, A., Yheskel, M., Kersjes, K., Li, J., Gatto, S., Takhar, M. et al. (2019). Discovery and preclinical evaluation of anti-miR-17 oligonucleotide RGLS4326 for the treatment of polycystic kidney disease. Nat Commun 10, 4148.

Liu, X., Vien, T., Duan, J., Sheu, S. H., DeCaen, P. G. and Clapham, D. E. (2018). Polycystin-2 is an essential ion channel subunit in the primary cilium of the renal collecting duct epithelium. Elife 7.

Ma, Y., Sugiura, R., Koike, A., Ebina, H., Sio, S. O. and Kuno, T. (2011). Transient receptor potential (TRP) and Cch1-Yam8 channels play key roles in the regulation of cytoplasmic Ca2+ in fission yeast. PLoS One 6, e22421.

Matsuo, Y., Asakawa, K., Toda, T. and Katayama, S. (2006). A rapid method for protein extraction from fission yeast. Biosci Biotechnol Biochem 70, 1992–4.

Mochizuki, T., Wu, G., Hayashi, T., Xenophontos, S. L., Veldhuisen, B., Saris, J. J., Reynolds, D. M., Cai, Y., Gabow, P. A., Pierides, A. et al. (1996). PKD2, a gene for polycystic kidney disease that encodes an integral membrane protein. Science 272, 1339–42.

Morris, Z., Sinha, D., Poddar, A., Morris, B. and Chen, Q. (2019). Fission yeast TRP channel Pkd2p localizes to the cleavage furrow and regulates cell separation during cytokinesis. Mol Biol Cell 30, 1791–1804.

Nakazawa, N., Xu, X., Arakawa, O. and Yanagida, M. (2019). Coordinated Roles of the Putative Ceramide-Conjugation Protein, Cwh43, and a Mn(2+)-Transporting, P-Type ATPase, Pmr1, in Fission Yeast. G3 (Bethesda) 9, 2667–2676.

Ong, A. C. and Harris, P. C. (2015). A polycystin-centric view of cyst formation and disease: the polycystins revisited. Kidney Int 88, 699–710.

Padhy, B., Xie, J., Wang, R., Lin, F. and Huang, C. L. (2022). Channel Function of Polycystin-2 in the Endoplasmic Reticulum Protects against Autosomal Dominant Polycystic Kidney Disease. J Am Soc Nephrol.

Palmer, C. P., Aydar, E. and Djamgoz, M. B. (2005). A microbial TRP-like polycystic-kidney-disease-related ion channel gene. Biochem J 387, 211–9.

Sato, M., Dhut, S. and Toda, T. (2005). New drug-resistant cassettes for gene disruption and epitope tagging in Schizosaccharomyces pombe. Yeast 22, 583–91.

Shen, P. S., Yang, X., DeCaen, P. G., Liu, X., Bulkley, D., Clapham, D. E. and Cao, E. (2016). The Structure of the Polycystic Kidney Disease Channel PKD2 in Lipid Nanodiscs. Cell 167, 763–773 e11.

Sinha, D., Ivan, D., Gibbs, E., Chetluru, M., Goss, J. and Chen, Q. (2022). Fission yeast polycystin Pkd2p promotes cell size expansion and antagonizes the Hippo-related SIN pathway. J Cell Sci 135.

Su, Q., Hu, F., Ge, X., Lei, J., Yu, S., Wang, T., Zhou, Q., Mei, C. and Shi, Y. (2018). Structure of the human PKD1-PKD2 complex. Science 361.

Torres, V. E. and Harris, P. C. (2014). Strategies targeting cAMP signaling in the treatment of polycystic kidney disease. J Am Soc Nephrol 25, 18–32.

Vien, T. N., Ng, L. C. T., Smith, J. M., Dong, K., Krappitz, M., Gainullin, V. G., Fedeles, S., Harris, P. C., Somlo, S. and DeCaen, P. G. (2020). Disrupting Polycystin-2 EF hand Ca(2+) affinity does not alter channel function or contribute to polycystic kidney disease. J Cell Sci.

Wilkes, M., Madej, M. G., Kreuter, L., Rhinow, D., Heinz, V., De Sanctis, S., Ruppel, S., Richter, R. M., Joos, F., Grieben, M. et al. (2017). Molecular insights into lipid-assisted Ca(2+) regulation of the TRP channel Polycystin-2. Nat Struct Mol Biol 24, 123–130.

